# Hierarchical confounder discovery in the experiment–machine learning cycle

**DOI:** 10.1101/2021.05.11.443616

**Authors:** Alex Rogozhnikov, Pavan Ramkumar, Rishi Bedi, Saul Kato, G. Sean Escola

## Abstract

The promise of using machine learning (ML) to extract scientific insights from high dimensional datasets is tempered by the frequent presence of confounding variables, and it behooves scientists to determine whether or not a model has extracted the desired information or instead may have fallen prey to bias. Due both to features of many natural phenomena and to practical constraints of experimental design, complex bioscience datasets tend to be organized in nested hierarchies which can obfuscate the origin of a confounding effect and obviate traditional methods of confounder amelioration. We propose a simple non-parametric statistical method called the Rank-to-Group (RTG) score that can identify hierarchical confounder effects in raw data and ML-derived data embeddings. We show that RTG scores correctly assign the effects of hierarchical confounders in cases where linear methods such as regression fail. In a large public biomedical image dataset, we discover unreported effects of experimental design. We then use RTG scores to discover cross-modal correlated variability in a complex multi-phenotypic biological dataset. This approach should be of general use in experiment–analysis cycles and to ensure confounder robustness in ML models.

## INTRODUCTION

The practice of training a model that maps high-dimensional input objects (such as biomedical images) to target labels (e.g. diseased vs. healthy) and subsequently quantifying model performance to identify biomarkers or disease phenotypes is of strong interest in multiple fields of diagnostic medicine (Wang et al. 2017; Rajpurkar et al. 2017; Gulshan et al. 2016; Ting et al.2017; Kermany et al. 2018; Zech et al. 2018). However, recent studies have shown that disease irrelevant features such as the physical site where an experiment is performed (Zech et al.2018) or the sample preparation protocol (Hägele et al. 2020) can severely bias these models. In these examples, “site” and “protocol” are potentially confounding discrete variables (**confounders**), whose values are additional **confounder labels** that accompany the target labels for each data point.

Many debiasing strategies exist to mitigate confounding effects on machine learning (ML) models including methods for (*i*) *a priori* balancing of datasets with respect to confounders (e.g. matching; see (Rosenbaum 2020), (*ii*) *post hoc* correction of datasets to reduce bias (e.g. restriction, stratification, harmonization, decorrelation; (Wallach and Heifets 2018; Hung 2019; Gerard and Stephens 2021), and (*iii*) incorporating bias resilience during model training (Zhang, Lemoine, and Mitchell 2018; Xu et al. 2018; Ganin and Lempitsky 2014; Zhao, Adeli, and Pohl 2020). However, while we have many tools to grapple with known confounders, we are lacking a general method to identify which variables in a set of potential confounders warrant debiasing.

A particular challenge is the attribution of bias to hierarchically organized **nested confounders** – confounders for which all data points that share a lower level confounder label also share the same higher level confounder label. For example, in induced pluripotent stem cell (iPSC) culture, multiple stem cell lines (“clones”) can be derived from the same human subject (“donor”) (Figure 1a). Donors differ due to genetic variation; clones are also known to differ though the biological underpinnings of clone-to-clone variability which remain an active area of research (D’Antonio et al. 2018; Carcamo-Orive et al. 2017; Volpato and Webber 2020). Such a confounder hierarchy is nested because each clone can only belong to a single donor. A typical linear approach to isolating the effect of a specific confounder is to compare models that fit the data with and without using the confounder of interest, i.e. by subtracting the variance explained by a model that excludes the confounder from the variance explained by a full model. However, for nested confounders this quantity will always be zero, because the lower level label (e.g., clone) confers perfect knowledge of the higher level label (e.g., donor), rendering this technique useless (Supplemental Figure S1a). Alternatively, one can assess for bias by computing the performance of linear decoders that predict confounder labels from data. This approach can work in the setting of nested confounders, but – unlike with calculations of variance explained – provides no concept of effect size: the prediction accuracy of two different confounders may be unmoored from the amount of structure they confer to the data (Supplemental Figure S1b). Beyond linear methods, graphical models are in theory purpose built for characterizing the effects of an interacting set of variables, including hierarchically organized ones, but they require assumptions about the data generating process as well as Bayesian inference methods which may not scale well to high dimensions (Chopin et al. 2015). Other nonlinear techniques like neural networks may be able to determine the effects of nested confounders, but they are sensitive to hyperparameter settings that may work for establishing the bias conferred by one confounder but not another, and thus cannot be deployed as a general method. One solution to dealing with a set of potential confounders is simply to apply debiasing strategies with respect to all such variables and test for improved model performance. However, if data is limited, model building expensive, or the relationships between confounders complex – and all three are often true in biology – this brute force approach is infeasible. Instead, a technique that does not suffer from the limitations articulated above and that can be used to quickly score the importance of potential confounders, including nested confounders, is needed.

**Figure 1.**
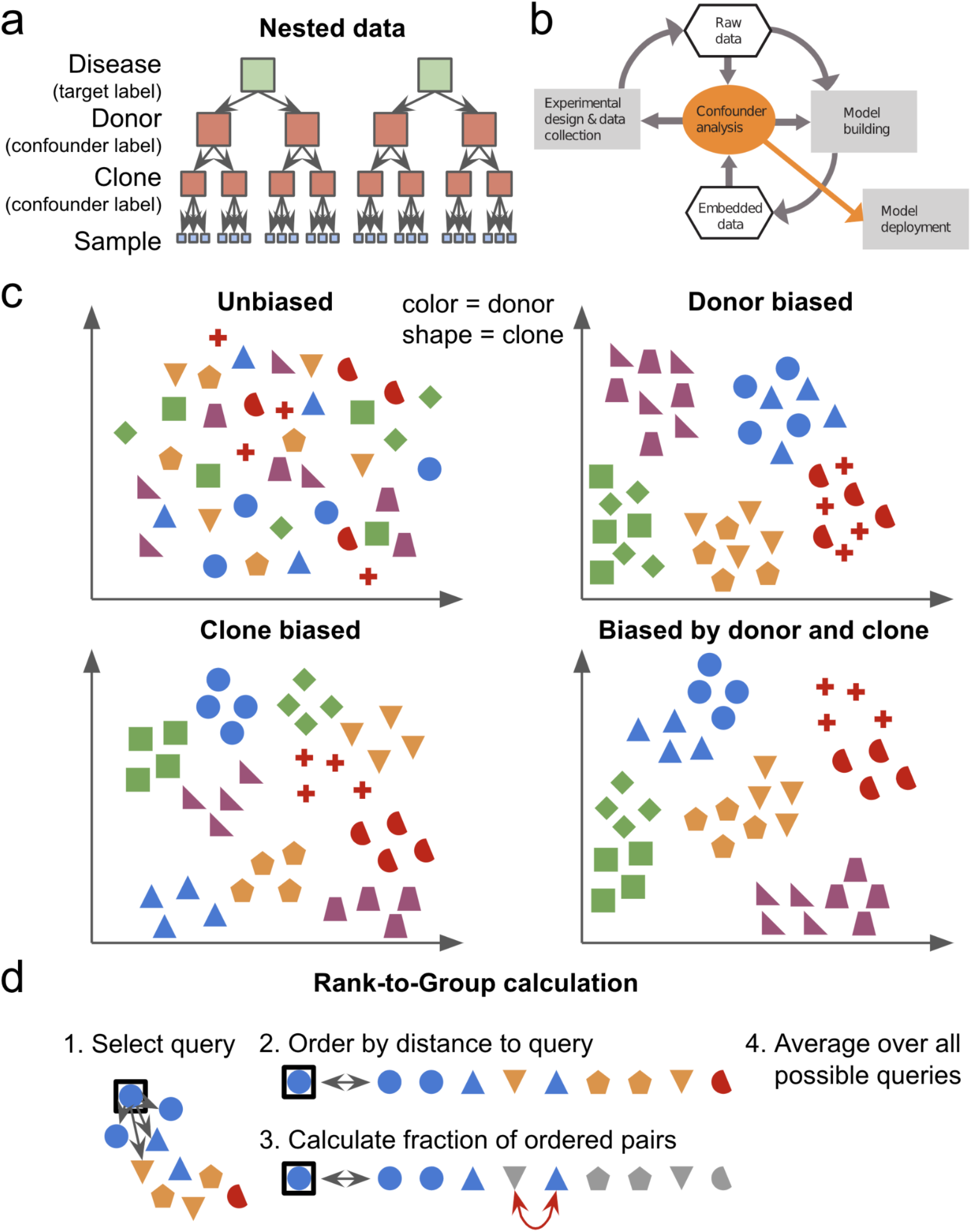
Hierarchical confounders and the RTG score. **a**. An example data hierarchy drawn from the field of stem cell biology. A typical modeling goal would be to predict the target label (healthy vs diseased) of a data sample. However, every data point would also be accompanied by potential confounder labels “clone” (i.e., stem cell line) and “donor” (i.e., human subject from which a clone is derived). Clone and donor are organized in a nested hierarchy: all data points that share the same clone label also share the same donor label. **b**. Role of confounder analysis in the iterative cycle of experimental data collection and model building with ML. The degree to which potential confounders confer structure to the raw data or data embeddings can inform both the experimental design used to collect more data and the modeling framework used to analyze the data. Model deployment should depend on confirmation of successful debiasing. **c**. Schematized data where donor is represented by color and clone by shape. The data is either unbiased by these variables (*top left*), biased by donor alone (*top right*), biased by clone alone (*bottom left*), or biased by both donor and clone (*bottom right*). Confounders can group the data. For example, for donor biased data (*top right*), the data are clustered by color, but within a cluster, no structure is conferred by shape. **d**. Diagrams illustrating the computation of the rank-to-group (RTG) score (Algorithm 1 in Methods) for donor (i.e., color). First, a query data point is selected and the distances from the query to all other data points are calculated (as schematized by the arrows in Diagram 1). Second, the data are ordered by their distances from the query (Diagram 2). Third, pairs of points are evaluated where one point shares the same donor as the query and the other does not. The RTG score for the current query is the fraction of such pairs for which the same donor point is closer to the query than the different donor point. In Diagram 3, one out of twenty possible pairs is out of order (the pair marked with the red arrow), giving an RTG score for this query of 0.95. The final RTG score is the average of each query-specific score over all possible query data points.

Consider Figure 1c which shows datasets with and without biasing by donor, clone, or both. Linear analysis would correctly declare that the data in the top right panel are biased by donor, but would also identify the data in the bottom left panel as donor dependent despite those data being constructed to be biased by clone alone. This is because the combined set of blue circles and triangles in this panel has substantially lower variance than the data as a whole as a result of the nested dependency between clone and donor. In order to correctly determine if donor confers structure beyond that determined by clone alone, we require a method that can disambiguate nested confounders. We could then use such a method to (1) inform the design of the next round of experiments to minimize bias, (2) guide data debiasing methods, or (3) build models that explicitly account for known biases (Figure 1b).

Here, we provide a novel non-parametric statistical method for scoring the degree to which data is structured by a potential confounder, the “Rank-to-Group” (RTG) score, which relies solely on similarity measures between data points. Thus, the RTG score is applicable both to raw data and to the embeddings that result from ML models. This method has a natural extension for handling nested confounders.

In the following sections, we describe the details of the RTG score method, provide analytic results for simple scenarios, compare it to linear analysis, and demonstrate how RTG scoring can be used to guide data collection and modeling decisions. We then apply our approach to two real-world datasets. First, we analyze a large public dataset of images of cultured cells (Cuccarese et al. 2020) and reveal that some features of the experimental design strongly bias the results. We furthermore show that linear techniques fail to discover these confounding effects. Next, we compare RTG analyses of a multi-modal dataset from patient-derived iPSC cultures (Shah et al. 2020) to interrogate the effects of donor, clone, and batch as well as their interactions. We show that these potential confounders differentially bias the data, but that their relative effects are conserved across three highly disparate modalities of biological measurement: quantitative PCR, brightfield microscopy, and single cell RNA sequencing. The general applicability of our approach to datasets with complex confounder hierarchies makes it of potentially broad utility when using ML techniques to interrogate large scale real world datasets.

## RESULTS

The **RTG score** is a non-parametric score of bias that is valid whenever (*i*) pairwise similarity scores between data points (e.g., distances) can be computed; and (*ii*) each data point has a set of associated features corresponding to potential confounders, which we term confounder labels. To compute the RTG score, first, for some “query” data point, all other points are considered in pairs where one member of each pair shares confounder label identity with the query point and the other does not. The **RTG query score** is the fraction of these pairs for which the same-confounder data point is more similar to the query than the different-confounder data point (Figure 1d and Algorithm 1 in Methods). Thus, this score measures the likelihood that a random data point that shares the same confounder label as the query is more similar to it than a random data point that does not. The average of the RTG query scores over all possible query points gives the final RTG score for the confounder in question. This score has a maximum of 1 when all data points that share the confounder label are closer to each other than they are to any other point. On the other hand, if the confounder confers no structure on the data, the RTG score will be 0.5 (with potential noise from finite size effects).

This score has several useful properties. First, it is well suited to high dimensional data because it depends on a similarity measure alone. Second, as a nonparametric rank order based score, it is insensitive to low noise levels that do not change the order of data similarities relative to the query, and, for the same reason, is largely insensitive to outliers. More generally, it provides useful results whenever distances are preserved locally even if they are less reliable over long length scales – as is the case for embeddings via UMAP, t-SNE, and other nonlinear techniques (Mclnnes et al. 2018) – since misorderings far from the query will likely have negligible impact on the calculated RTG score. Third, it can be applied to any embedding, regardless of interpretability. Finally and most importantly, this technique is easily extended to the case of nested confounders.

When confounders are hierarchically nested (e.g., donor and clone) the RTG score for the higher level will include effects of the lower level, obscuring the effects of the higher level alone. To disambiguate the effects of two hierarchically related confounders, we can compute a **restricted RTG score**. For example, during the evaluation of the “donor-exclude-same-clone” RTG score for a specific query, we restrict the computation to exclude all data points that share the same clone label as the query (Supplemental Figure S2 and Algorithm 1 in Methods). This means that the RTG query score measures the degree to which, for data points that do not share the same clone label as the query, similarity with the query can sort those data into same-donor and different-donor groups. If an apparent confounding effect of donor is entirely attributable to clone as in the bottom left panel of Figure 1c, the donor-exclude-same-clone restricted RTG score will be at the chance level of 0.5. Thus, the restricted RTG score allows us to disambiguate clone-confounder effects from (donor-and-clone)-confounder effects (bottom left and right panels of Figure 1c).

While we only consider examples of discrete valued confounding variables in this article, the RTG and restricted RTG scores can be extended to the setting of continuous valued confounders as described in the Appendix.

### Analytic RTG scores for hierarchical Gaussian mixture data

We now illustrate how RTG scores can disambiguate between three scenarios of confounder-introduced variability by applying them to synthetic data. We specify a three-level clone–donor–sample hierarchical Gaussian mixture model governed by three parameters, the variances, 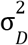, 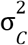, 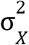 (Figure 2a). The relative sizes of these variances define the structure of the data. We consider three extreme scenarios: data fully biased by both donor and clone, donor only, or clone only (Figure 2b). For these cases, we can compute simple analytic expressions for the RTG scores of clone and donor and for the restricted RTG score of donor-exclude-same-clone. These depend only on *D* and *C*, the number of donors and clones per donor respectively, assuming large numbers of samples per clone (Figure 2c). If we interpolate between the donor-and-clone biased scenario and the donor only and clone only scenarios, we observe that the RTG scores indeed change according to the analytic expressions (Figure 2d).

**Figure 2.**
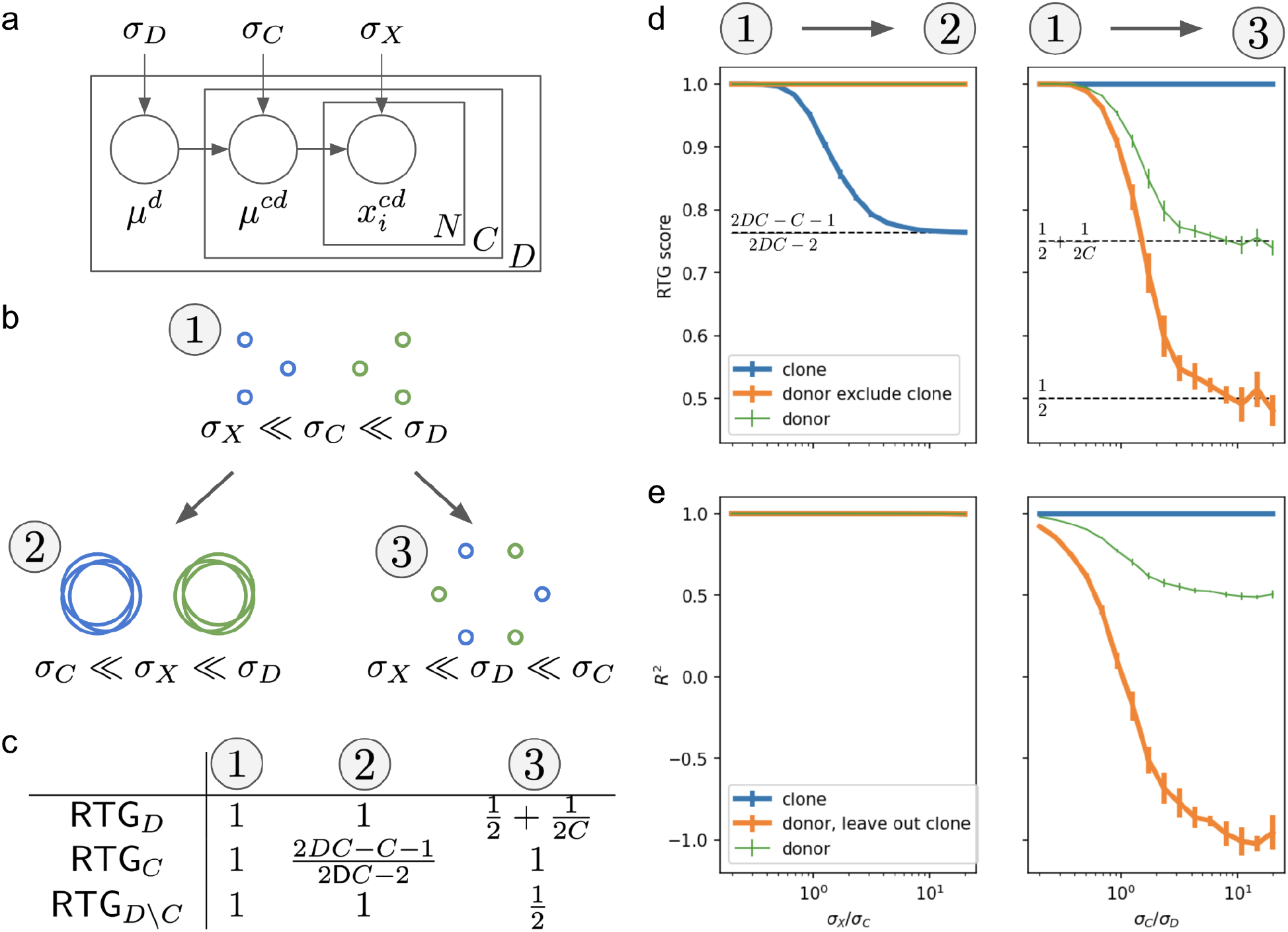
Confounder analysis of hierarchical Gaussian synthetic data. **a**. Data generation scheme. The means for each donor and clone are sampled as 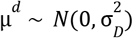 and 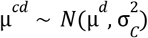 respectively. Then, the *i*^th^ data point for each clone is sampled as 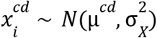. *D, C*, and *N* are the number of donors, clones per donor, and data points per clone. **b**. Three extreme confounder scenarios for 2D data. Each circle represents a single clone whose radius represents σ_*X*_. Each color represents a donor. In scenario 1, the variance differentiating donors dominates the variance differentiating the clones within a donor which in turn dominates the variance of the data for each clone (i.e., 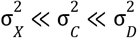). Thus, the data are completely biased by both donor and clone. With variances ordered as 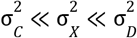 (scenario 2) and 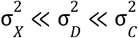 (scenario 3), the data are instead biased by donor alone or clone alone respectively. **c**. The analytic RTG scores for donor, clone, and donor-exclude-same-clone for the extreme scenarios 1, 2, and 3 from **b**. **d**. The evolution of RTG scores as datasets change from scenario 1 to scenarios 2 (*left*) and 3 (*right*). RTG scores are plotted against the ratios σ_*X*_/σ_*C*_ (*left*) and σ_*C*_/σ_*D*_ (*right*) while ensuring the other constraints from **b** (i.e., that σ_*D*_ is large for the left panel and σ_*X*_ is small for the right panel) remain in place. RTG for donor (green) is plotted with a thin line since it does not isolate the confounding effects of a single potential confounder as opposed to donor-exclude-same-clone (orange) which does. **e**. Leave-one-out cross validated *R*^2^ values of linear models fit using clone and donor identities, and – to isolate donor effects – the leave-clone-out cross validated *R*^2^ values for models fit using donor but only trained on *C* – 1 clones and tested on the held out clone. Other conventions as in **d**. Left panels in **d** & **e**: *D* = 2; *C* = 10; σ_*D*_ = 400σ_*C*_. Right panels: *D* = 50; *C* = 2; σ_*X*_ = 10^−4^ σ_*D*_. All panels: *N* = 100; data is 10 dimensional; error bars are std devs over 5 random samples of the data.

Our analytic results illustrate the importance of restricted RTG scores for distinguishing between single or multiple confounder effects. For example, for data biased only by clone (i.e., when 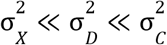), the analytic expression for the RTG score for donor (denoted *RTG_D_*) is 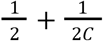, which for small *C* can significantly exceed 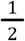 even though in truth there is no donor effect (right panel of Figure 2d). However, while *RTG_D_* be may be misleading, donor-exclude-same-clone restricted RTG (denoted *RTG_D\C_*) reports a score of 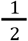, consistent with no donor effect. If, on the other hand, donor does structure the data beyond the effects of clone (i.e., when 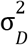 is similar to or greater than 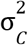), then *RTG_D\C_* will exceed 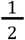, confirming the effect.

### Linear-model confounder analysis approaches fail on hierarchical data

Confounder discovery via linear analysis is a reasonable baseline against which to compare the RTG method. However, as discussed in the Introduction, the standard approach of comparing full and partial models to isolate the effect of a potential confounder – i.e., by computing the difference in data variance explained by models that use and do not use the confounder of interest – fails for nested hierarchies (see Supplemental Figure 1a). An alternative linear approach that avoids this pitfall is “leave-confounder-out” cross validation. To isolate the confounding effects of a variable higher in a confounder hierarchy like donor, a model can be constructed from donor labels using only a subset of the data that holds out one or more clones. This model can then be tested, in terms of the variance explained, on the held out data. This approach is similar to K-fold cross validation except that the data folds, rather than being random, are defined by the lower level label identities (e.g., clone).

Comparison of the left panels of Figures 2d–e shows the utility of RTG scoring versus linear analysis. Data that are biased by both donor and clone or by donor alone are indistinguishable by *R*^2^ values but clearly differentiated when using the RTG method. In contrast, leave-confounder-out linear analysis is useful for disambiguating clone biased data from data biased by both donor and clone (Figures 2d–e, right). Note that if the number of labels for a held out lower level confounder (e.g., clone) is two and if the structure of the data is dominated by that lower lever confounder, then the leave-confounder-out *R*^2^ for the higher level variable will reach a minimum of −1 for hierarchical Gaussian data (as seen for the orange line in Figure 1e, right).

We next consider the consequences of noise in synthetic data with mixed donor and clone effects (Figure 3). For increasing noise values, RTG scores are largely stable while *R*^2^ values drop precipitously. This is because RTG scoring – a rank order based method – is unaffected by noise unless distances to the query of data points that share and do not share confounder labels change order (see Figure 1d and Algorithm 1 in Methods). In other words, as long as noise does not result in mixing of clusters in the data that are defined by confounder labels, RTG analysis will reveal the confounder dependent structure in the data. *R*^2^ values, on the other hand, only report the structure of the data indirectly through their measurement of explained variance, and thus are guaranteed to drop as data noise increases even when the underlying clustered structure in the data remains intact.

**Figure 3.**
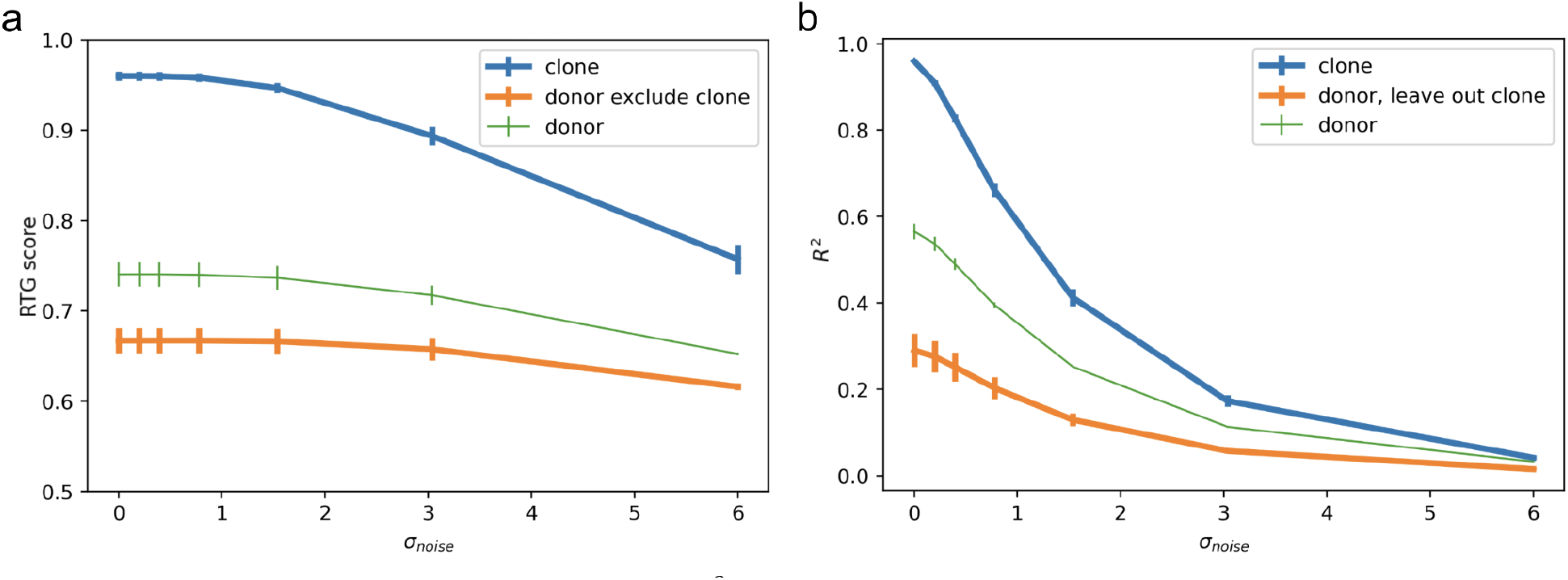
Comparison between RTG scores (**a**) and *R*^2^ values (**b**) as a function of noise. The data is generated as per Figure 2a, but then randomly projected to a much higher dimensional space before being corrupted by white noise with scale σ_*noise*_. Conventions are as in Figures 2d–e except: *D* = 14; *C* = 2−6 (different for each donor); σ_*D*_ = σ_*c*_ = 1; σ_*X*_ = 0.25; *N* = 25; data is 2 dimensional before being projected to 512 dimensions; error bars are std devs over 3 random samples of the data.

### Model debiasing example

We can consider a simple example of using RTG analysis to guide decision making around data collection and modeling when building classifiers to predict disease state (as per Figure 1b). We first generate synthetic data from a mixture-model according to Figure 2a, but with the modification that half of all donors – representing healthy human subjects – have the vector [1,0,…, 0]^*T*^ added to their means. The other half of the donors have their means shifted by [−1,0,…, 0]^*T*^ and represent diseased patients. As before, the confounder structure of the generated data depends on the relative values of the variances of donor, clone given donor, and data sample given clone (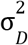, 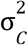, and 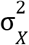).

First, we consider the case of analyzing small datasets – two donors per disease state and two clones per donor – to determine whether or not more donors or clones should be collected to improve the disease state prediction accuracy of a logistic regression model. When the data is biased by donor but not clone, adding more donors while keeping the number of clones per donor fixed improves model performance while adding clones confers no benefit (Figure 4a). Conversely, for data biased by clone but not donor, adding either clones or donors improves model performance equally (Figure 4b). These results demonstrate the power of RTG analysis for informing data collection strategy. In a real world setting, new samples of lower level variables (e.g., more clones) may be cheaper to obtain than samples of higher level variables (e.g., donors). The confounder structure as measured by RTG scoring can dictate how best to use resources for data collection by clarifying when sampling more higher level confounders is necessary or when adding new lower level examples to a dataset is sufficient.

**Figure 4.**
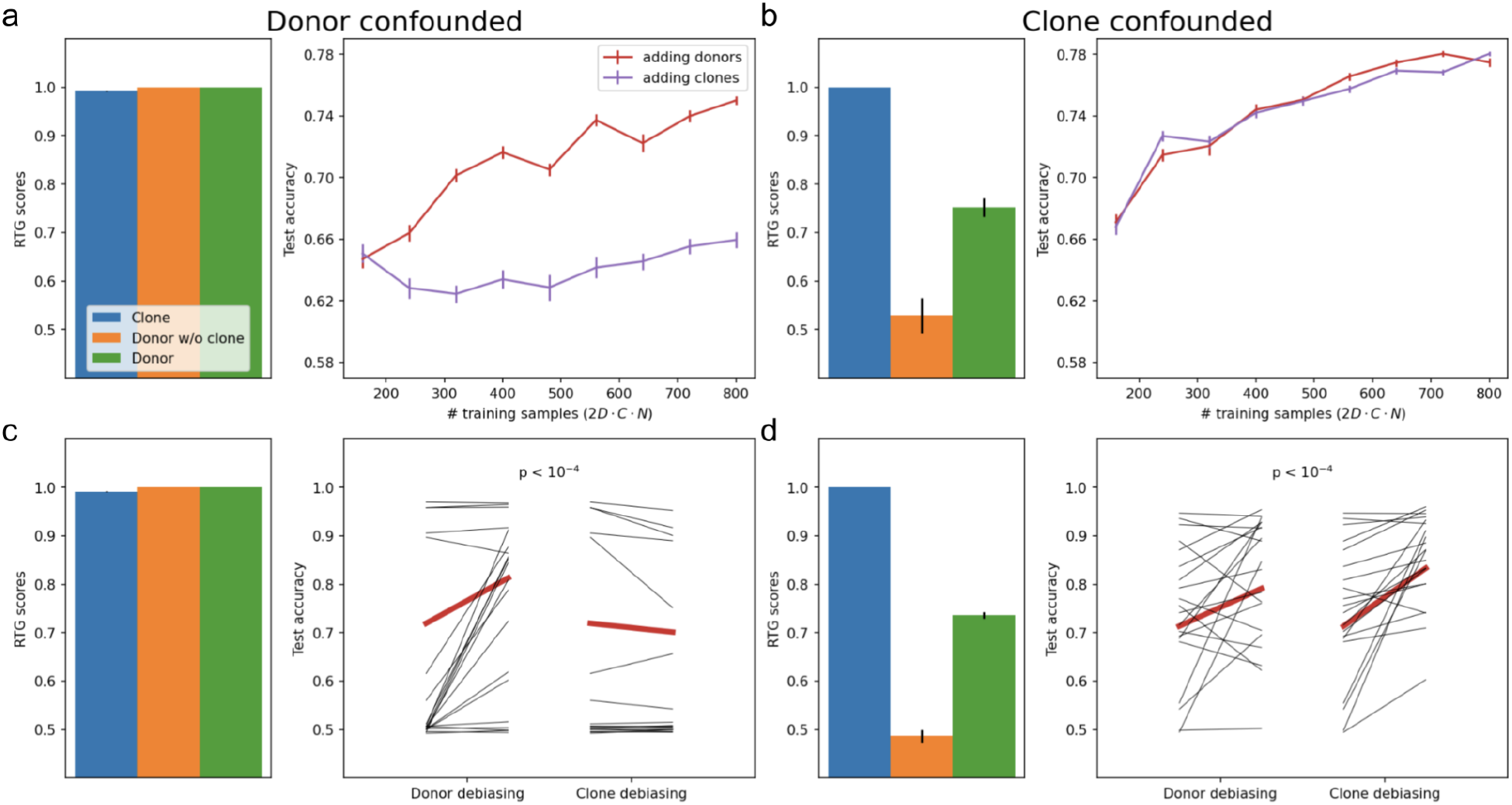
Using RTG scores to inform data collection and debiasing. **a**. Effect of training data quantity on model performance for donor confounded data. Left: RTG scores for donor, clone, and donor-exclude-same-clone before adding new donors or clones. Right: Accuracy of logistic regression models on test data. Models are trained while either increasing the number of donors in the training dataset when holding the number of clones per donor fixed (*D* = 2−10 and *C* = 2, red) or while increasing the number of clones per donor when holding the number of donors fixed (*D* = 2 and *C* = 2−10, purple). **b**. Same as **a** for clone confounded data. **c**. Effect of data debiasing on model performance for donor confounded data. Left: RTG scores prior to debiasing. Right: Changes in model performance when debiasing by donor or by clone. Debiasing by donor occurred as follows. For each donor, the centroid of the corresponding disease state was subtracted from that donor’s centroid to obtain that donor’s specific direction. PCA was performed on the set of donor specific directions and the top two PCs were identified as the donor specific subspace. The data was then projected out of that subspace. Debiasing by clone was identical except that each clone’s specific direction was computed from the difference between that clone’s centroid and the centroid of the corresponding donor. Red lines show mean changes due to debiasing over 100 sample datasets; black lines show 20 example datasets. The mean change due to donor debiasing was significantly different from the mean change due to clone debiasing (p < 10^−4^ via resampling). **d.** Same as **c** for clone confounded data. **a**: σ_*D*_ = 1; σ_*C*_ = 0.1. **b**: σ_*D*_ = 0.1; σ_*C*_ = 1. **a** &**b**: σ_*X*_ = 0.1; *N* = 10 for training; 10000 test data points each with a unique donor and clone; 20 dimensional data; error bars are s.e.m. over 20 random samples of the training data. **c**: σ_*D*_ = σ_*C*_ = σ_*X*_ = 0.3 in all dimensions except 3 dimensions with σ_*D*_ = 100; *D* = 4, *N* = 10. **d**: σ_*D*_ = σ_*C*_ = σ_*X*_ = 0.3 except 3 dimensions with σ_*C*_ = 100; *D* = 2, *N* = 20. **c** & **d**: *C* = 2; 10000 test data points; 10 dimensional data.

Next, we consider the case of using RTG analysis to inform how best to improve model quality when no further data collection is possible. For this example, during data generation, we confine 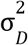 and 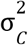 to only be large in a subset of data dimensions when biasing by donor and clone respectively. Then, to debias with respect to donor, for example, we determine directly from the data which dimensions correspond most strongly to donor and project the data out of those dimensions prior to model fitting. For donor biased data, average prediction accuracy on test data improved from 72% before debiasing by donor to 81% afterwards, while debiasing by clone showed a small negative effect (70% post-debiasing accuracy; Figure 4c). The difference between these mean changes was highly significant (p < 10^−4^ by resampling; Supplemental Figure S3a). For clone biased data, average prediction accuracy improved from 71% before debiasing by clone to 83% afterwards, while debiasing by donor showed a smaller, though meaningful effect (79% post-debiasing accuracy; Figure 4d). The difference between these mean changes was also highly significant (p < 10^−4^; Supplemental Figure S3b). These results demonstrate that in the data constrained regime, identification of confounders with large effects can inform strategies for building higher performing models.

### Identification of confounding effects in a hierarchical biomedical dataset

Next, we applied RTG analysis to identify confounders in a dataset of biomedical image embeddings released in the public domain by Recursion Pharmaceuticals (Cuccarese et al.2020). The raw images are of cell cultures in 1536-well plates. The data consists of 2 experimental groups of 12 plates each. Each well is imaged at four sites. The wells along the edges of the plates were not used. Many of the wells in this dataset were treated with pharmacologic agents, but for our analysis, we chose to look only at untreated control wells, which were randomly dispersed across each plate. This left a dataset of approximately 8,000 image embeddings that we analyzed for the confounding effects of experimental group ID (1 or 2), plate ID (1 to 12), well position (e.g., C4), unique well ID (e.g., C4 on plate 3 in experiment 1), camera field of view within a well or “imaging site” (1 to 4), and whether or not an image came from a well on the “subedge” – the outer rim of wells that remain after excluding the edge wells themselves.

In Figure 5a we calculated RTG scores and restricted RTG scores for these potential confounders, and confirmed via subsampling that our calculated scores are precise enough to facilitate comparison between them (Supplemental Figure S4). Our analysis reveals that the Recursion data is largely free of obvious experimental pitfalls. For example, imaging site does not bias the data (RTG 0.5) while unique well ID nearly completely biases the data (RTG 0.95), meaning that the four images within each well are essentially interchangeable. Furthermore, well position alone strongly biases the data (RTG 0.76), but well-position-exclude-same-plate (i.e., equivalent to well-position-exclude-same-unique-well) drops the RTG score to 0.58, supporting the notion that similarity between images within the same well primarily drives the embedded data structure. Next, we observe that the confounding effects of plate ID (RTG 0.55) can be entirely explained by experiment ID (plate-exclude-same-experiment RTG 0.5). Experiment ID, on the other hand, confers significant structure to the data (RTG 0.87), suggesting some difference in conditions between experimental groups. Each experiment was a different “batch” of data (Cuccarese et al. 2020), suggesting the presence of confounding effects by batch.

**Figure 5:**
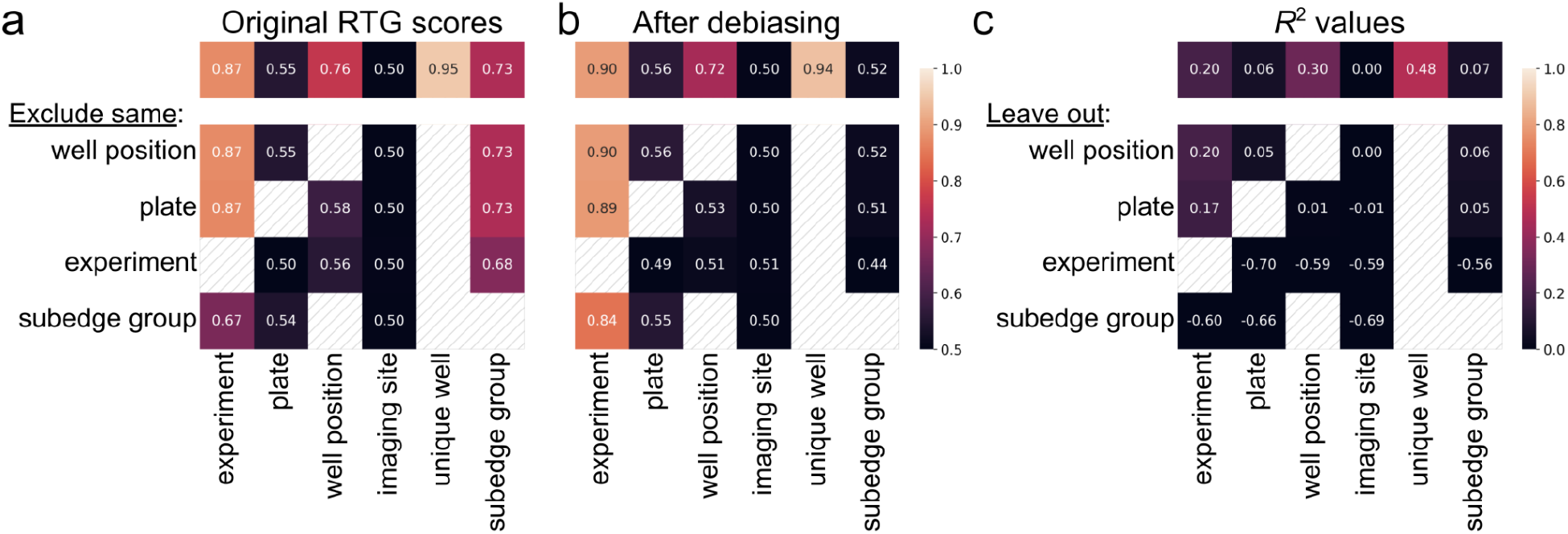
RTG scores and *R*^2^ values calculated from the analysis of Recursion Pharmaceuticals’ public dataset (Cuccarese et al. 2020). **a**. RTG scores for potential confounders: experiment, plate, well position, imaging site, unique well, and whether or not a data point is from a subedge well. Each confounder is considered with (top row) and without (other rows) the exclusion of each other confounder (except in the setting where the would-be excluded confounder is strictly higher in the confounder hierarchy, making exclusion impossible). **b**. RTG scores after the data has been collapsed along the dimension connecting the means of the subedge well data and the non-subedge well data. Conventions as in **a**. **c.** Five-fold cross validated *R*^2^ values (top row) and leave-confounder-out cross validated *R*^2^ values (other rows) for the same potential confounders as in **a**. Negative values mean that models built while leaving out data with a particular confounder label and then tested on that left-out data are worse than a naïve model that simply predicts the mean of the whole dataset.

However, we found that whether not an image comes from a well on the subedge strongly structures the data (RTG 0.73). Indeed, subedge membership is a strong influence even across experiments (subedge-exclude-same-experiment RTG 0.68). These results suggest that despite not using the edge wells – presumably to mitigate a common issue with well plates – the data embeddings remain highly influenced by whether or not a well is on the outer rim of the in-use wells. To confirm this finding, we made a simple manipulation of the data by removing the dimension defined by the line that connects the centroids of the subedge well data and the non-subedge well data. This debiased data showed improved RTG scores (Figure 5b). The residual effect of well-position-exclude-same-plate is largely eliminated (RTG from 0.58 to 0.53) with the remaining confounding effect attributable to experiment (well-position-exclude-same-experiment RTG 0.51). The confounding effects of experiment ID become slightly more pronounced (RTG from 0.87 to 0.9) including when excluding the effects of subedge membership (experiment-exclude-same-subedge-group RTG from 0.67 to 0.84), suggesting that simple manipulations of the data can mitigate certain confounding effects while amplifying the effects of other confounders.

We can compare the results of RTG and linear analysis to demonstrate the power of the RTG approach (Figures 5a and 5c). Though some of the conclusions are the same (e.g., that images in the same well are more correlated than any other grouping), others differ. For example, well position accounts for more variance than experiment though, per RTG analysis, experiment confers more clustered structure to the data. Furthermore, the highest *R*^2^ values only reach about half of their maximum while the highest RTG scores are nearly maxed out. RTG analysis specifically measures the degree to which a confounder clusters data while *R*^2^ captures this only indirectly through a measurement of variance explained. Thus, the Recursion data demonstrates that real world data embeddings can be both highly clustered and noisy, suggesting the risk of a faulty conclusion if relying on a low *R*^2^ value as a surrogate for low bias.

Most importantly, linear analysis fails to unambiguously identify the confounding effects of the subedge. The raw cross validated *R*^2^ for subedge is near zero (0.07, last value in first row of Figure 5c) which is in conflict with the negative *R*^2^ values obtained when performing leave-subedge-group-out cross validation (bottom row). These negative values suggest that a model fit with data from the subedge is poor at predicting the data off the subedge and vice versa (à la the negative values in the right panel of Figure 2e). Such conflicting results may impede discovery of the importance of subedge if using linear confounder analysis. Of note, leave-experiment-out cross validation also results in negative values (next to bottom row), but, unlike for subedge, the raw cross validated *R*^2^ for experiment is well above zero (0.20, first value in first row).

### RTG analysis yields modality independent confounder effects in multi-phenotypic data

Next, we applied RTG analysis to our own multi-phenotypic dataset collected from human iPSC-derived brain organoids (Shah et al. 2020). This data consists of three kinds of measurements: gene expression via quantitative polymerase chain reaction (qPCR), morphology via brightfield microscopy, and cell type distribution via single cell RNA sequencing. For each modality, multiple donors and clones per donor were used and data collection occurred in several experimental batches. We analyzed 21 donors, 58 clones, and 17 batches for qPCR; 10 donors, 20 clones, 7 batches for brightfield; and 14 donors, 29 clones, and 3 batches for scRNAseq.

Our analysis (Figure 6) detects substantial confounder effects. Both clone RTG and donor-exclude-same-clone restricted RTG are elevated with clone showing a somewhat stronger effect than donor. Batch shows a weak effect on its own, but a confounder defined as the intersection of batch and clone (i.e., where each confounder label is the union of a batch label and a clone label) strongly biases the data. Similar, though weaker, confounding effects are seen with the intersection of batch and donor. These results illustrate the utility of RTG scoring in the cycle of experimentation and data analysis (Figure 1b). They suggest that future data collection can be targeted at increasing the number of clones which is typically less resource intensive than increasing donors. Furthermore, both clones and donors should be distributed across batches to reduce the bias conferred by these variables. Finally, these results clarify the need to use ML techniques that can generate data embeddings which are insensitive to donor, clone, and batch effects in order to aid biological interpretation.

**Figure 6.**
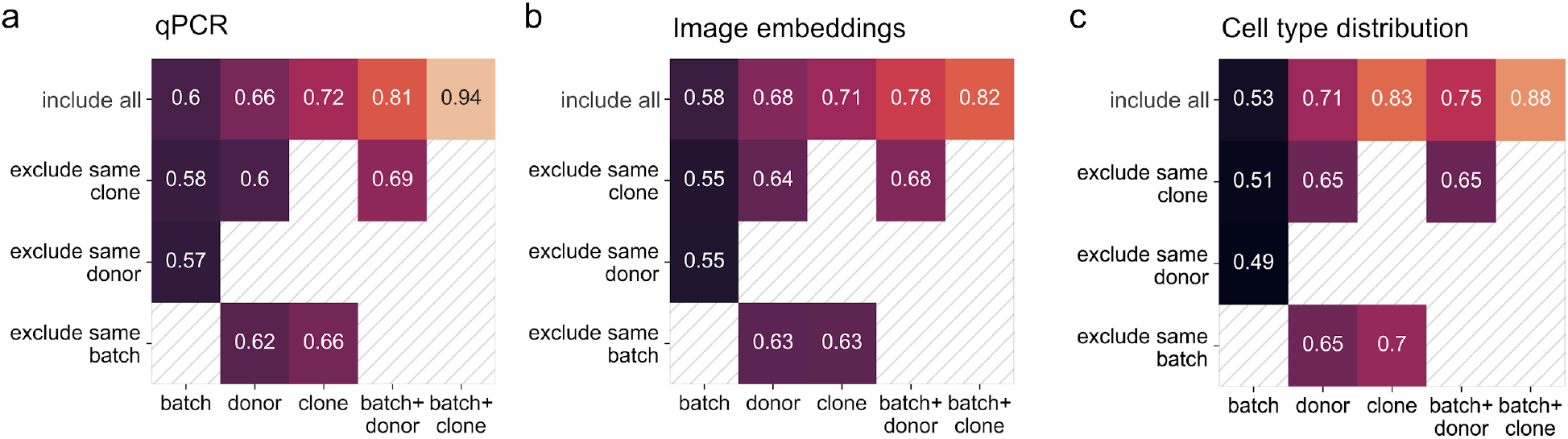
RTG scores when evaluating different aspects of our previously published multimodal dataset (Shah et al.2020). For each modality, potential confounders batch, donor, and clone are evaluated, as well as the intersections of batch plus donor and batch plus clone. Restricted RTG scores excluding batch, donor, and clone are also presented (unless they are higher in the data hierarchy than the confounder being evaluated). **a**, **b**, and **c** show RTG scores computed from gene transcriptional data via qPCR, image data via brightfield microscopy, and cell type distribution data via scRNAseq for iPSC-derived brain organoid cultures.

Strikingly, we see that the RTG scores for gene expression, imaging, and cell type distribution (Figures 6a–c) are consistent across modalities (average pairwise Spearman’s rank correlation of 0.924). These results suggest that there is a common biological origin of these phenotypic modalities; thus, the use of cost-efficient, scalable modalities (e.g., imaging) for the characterization of variability may be sufficient without the need to scale-up highly resource intensive techniques such as single cell sequencing (Shah et al. 2020). This demonstrates another use for confounder analysis in the experiment–analysis cycle: in addition to informing how to balance future data collection (e.g., more clones), RTG scores can also inform resource allocation toward specific types of assays.

## DISCUSSION

Strategies for confounder analysis are currently lacking. Standard linear methods suffer from outlier sensitivity, are blind to complex structure in data, and cannot disambiguate hierarchically nested confounders; matching and stratification strategies suffer on high-dimensional data due to combinatorial scaling of unmatched dimensions, and matching is impossible for a lower level confounder in a nested hierarchy; and Bayesian models suffer from poor performance in high dimensions (Chopin et al. 2015). We presented a novel non-parametric method, the rank-to-group or RTG score that addresses these issues and is easily deployable in settings where data interpretability is limited (e.g., neural network embeddings) and confounder sensitivity of great concern (e.g., models of biomedical data that are intended to support disease diagnosis or treatment). The only requirements of our method are (*i*) ability to compute a similarity score between data points and (*ii*) knowledge of confounder labels for each data point. We have demonstrated that RTG scoring is robust to noise and can identify the level in a confounder hierarchy from which a confounding effect arises even when the structure conferred by the confounder is nonlinear.

When we apply our method to real world biomedical datasets, we find that it can identify confounders such as batch, edge, donor, and clone, a point of critical importance when attempting to derive general results from these kinds of data. Batch effects – which arise when some experimental conditions shift despite the intention of repeating them exactly – is an example of a confounder that iterative experimental design and careful data collection may be able to mitigate. Donor effects may be reduced by matching certain variables across disease and healthy groups (e.g., gender and age) but may also require strict protocols (e.g., sample collection and handling) to mitigate fully. Edge effects can be partially managed by randomization and clone effects by increasing the numbers of clones per donor, but each of these confounders also likely requires specific computational solutions to mitigate. Recent methods that force data embeddings to be insensitive to certain potential confounders (Ganin and Lempitsky 2014; Zhao, Adeli, and Pohl 2020) may be of particular value for debiasing data with respect to confounders such as edge and clone. However, they rely on minimizing the decodability of a confounder in data embeddings which does not address the importance of that confounder in structuring the data (i.e., one confounder may be more decodable than another while structuring the data more weakly; see Supplemental Figure S1b). In all of these cases, a method such as RTG scoring is needed to identify the critical confounders and thus inform experimental and data analysis choices that can ultimately improve models of the data.

One of the shortcomings of RTG scoring is that, in our implementation, we have not proposed a method for estimating confidence intervals. Though our subsampling results of the Recursion Pharmaceutical data (Supplemental Figure S4; Cuccarese et al., 2020) reassure us of the stability of the RTG method, at least for that particular dataset, it is not a rigorous result. However, this could be addressed by implementing bootstrapping techniques that respect the data hierarchy (Saravanan, Berman, and Sober 2020).

RTG scoring is more than just a *post hoc* tool for comparing whether one data embedding is better – more confounder resilient – than another. Rather, we envision that the utility of our approach will be best realized as part of a virtuous cycle of experimental design, data collection, modeling building, and confounder analysis. Careful attribution of confounding effects can give confidence as to an ML model’s likely performance after deployment. If, as in the example used throughout this article, a model’s ability to identify disease state is simply due to the fact that it has separated every clone in embedding space, there may be considerable concern that this model has not learned anything about disease per se. In this case, RTG analysis can can guide how best to improve confounder robustness through both new experiments (i.e., by suggesting the highest value new data to collect) and updated model training approaches (i.e., that are specifically designed to counter the effects of certain confounders) so as to mitigate confounder influences in the next cycle of development. The general applicability of this approach to high-dimensional datasets with complex, potentially nested, confounder hierarchies makes it of broad utility when using ML techniques to interrogate large scale real world datasets.

## METHODS

The Rank-to-group score algorithm is articulated in Algorithm 1. Additionally, Python code is available at https://github.com/herophilus/rtg_score.

### Algorithm 1: Rank-to-group score

We start with (*i*) the items in our dataset *x_i_* for *i* = 1,…, *N*, (*ii*) an included group variable *I* whose confounding effect we want to estimate, and (*iii*) zero or more excluded group variables *E*^1^,…, *E^L^* whose confounding efforts we wish to ensure are not misattributed to *I*. Each item *i* has a label for the included group of interest *I_i_* and – optionally, when excluding other potential confounders – labels for the excluded groups 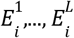.

We compute *RTG_q_*, the RTG score for query item *q*, as follows:

1. Define two sets: *S_q_* consisting of the indices of the items that share the same included group label *I_q_* with item *q*, and *D_q_* consisting of the indices of the items whose included group labels do not equal *I_q_*. When excluding other potential confounders, the indices of the items that share the same excluded group labels 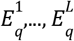 with item *q* are removed from *S_q_* and *D_q_*.

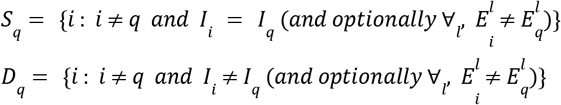
2. Then, the *RTG_q_* is defined as the fraction of pairs of items from indices in sets *S_q_* and *D_q_* where the query item is more similar to the item whose index is in set *S_q_* than it is to the item whose index is in set *D_q_* (with half attribution in the case of equal similarities):

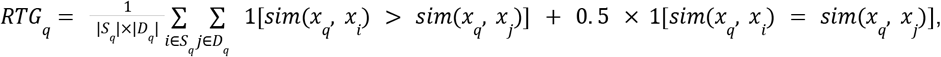

where |*S_q_*| and |*D_q_*| are the number of elements in the sets *S_q_* and *D_q_*. This definition is identical to the area under the curve (AUC) of the receiver–operator characteristic (ROC) curve computed when (*i*) all the items whose indices are in the set *S_q_* ∪ *D_q_* are sorted by their similarity to the query item and labeled with 1s and 0s if their indices are members of *S_q_* and *D_q_* respectively, and (*ii*) the 1s and 0s are treated as true and false positives as in standard ROC analysis. Thus we can efficiently compute *RTG_i_* from a single pass through *S_q_* ∪ *D_q_* after sorting.

The final RTG score is simply the average of the scores *RTG_q_* over all possible query items in the dataset (i.e., *q* = 1,…, *N*). Of note, for some queries, *RTG_q_* may be undefined because either set or *S_q_* or *D_q_* is empty. These queries are excluded from averaging.

## ACKNOWLEDGEMENTS

The authors would like to thank S Linderman and L Abbott for helpful discussions and comments on the manuscript.

## AUTHOR CONTRIBUTIONS

AR, RB: conceptualization; AR and GSE: simulations; AR: data analysis; GSE: analytics; PR and GSE: writing—original draft; SK and GSE: writing—review and editing

## APPENDIX

### RTG scoring with continuous confounders

The RTG score defined in Algorithm 1 assumes that potential confounders are discrete (e.g., gender). However, in many situations confounders can be continuous (e.g., age, or some companion measurement in an experiment such as temperature). In this case, RTG scores cannot be calculated as currently defined. One simple option is to discretize continuous confounders into bins. In this case, one needs to choose the bin boundaries which represent hyperparameters of the RTG score.

Alternatively, we can relax the definition of the RTG score to explicitly permit continuous confounders. Using notation from Algorithm 1, we have:

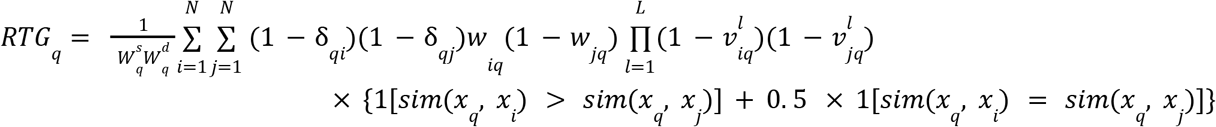

where: 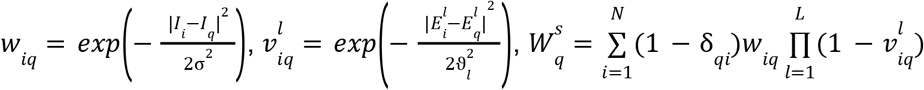, and 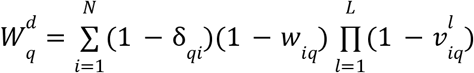. Here, *I_i_* and 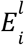 are the values of continuous included and excluded confounders for data point *i*; *w_iq_* and 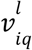 measure, from 0 to 1, how similar an included or excluded confounder value is to the corresponding confounder of query item *q*; δ_*qi*_ is the Kronecker delta; and 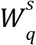 and 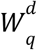 are normalizers that “count” the number of items that share and do not share included confounder identity with the query item after items that share excluded confounder identity with the query have been removed.

If, for discrete confounders with labels rather than values, we define |*I_i_* − *I_q_*| = 0 for *I_i_* = *I_q_* and |*I_i_* − *I_q_*| = ∞ for *I_i_* ≠ *I_q_*, then this equation for *RTG_q_* is identical to the equation in Algorithm 1.

Thus it is possible to mix and match continuously-valued and discrete valued confounders.

When dealing directly with continuous confounders, one needs to choose the hyperparameters σ and ϑ_*l*_ which determine the scale over which confounder variables can be said to switch from being of the same value to being of different values. These hyperparameters are analogous to the bin boundaries that need to be chosen when using the discretization strategy. Thus, either strategy requires determination of hyperparameters which is unnecessary for purely discrete confounders. Of note, it is appropriate to to call these hyperparameters rather than parameters because they relate to the confounders *I_i_* and 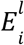 and not to the data *x_i_*. RTG analysis applied to data with continuous valued confounders remains nonparametric with respect to the data itself.

### SUPPLEMENTAL FIGURES

**Supplemental Figure S1.**
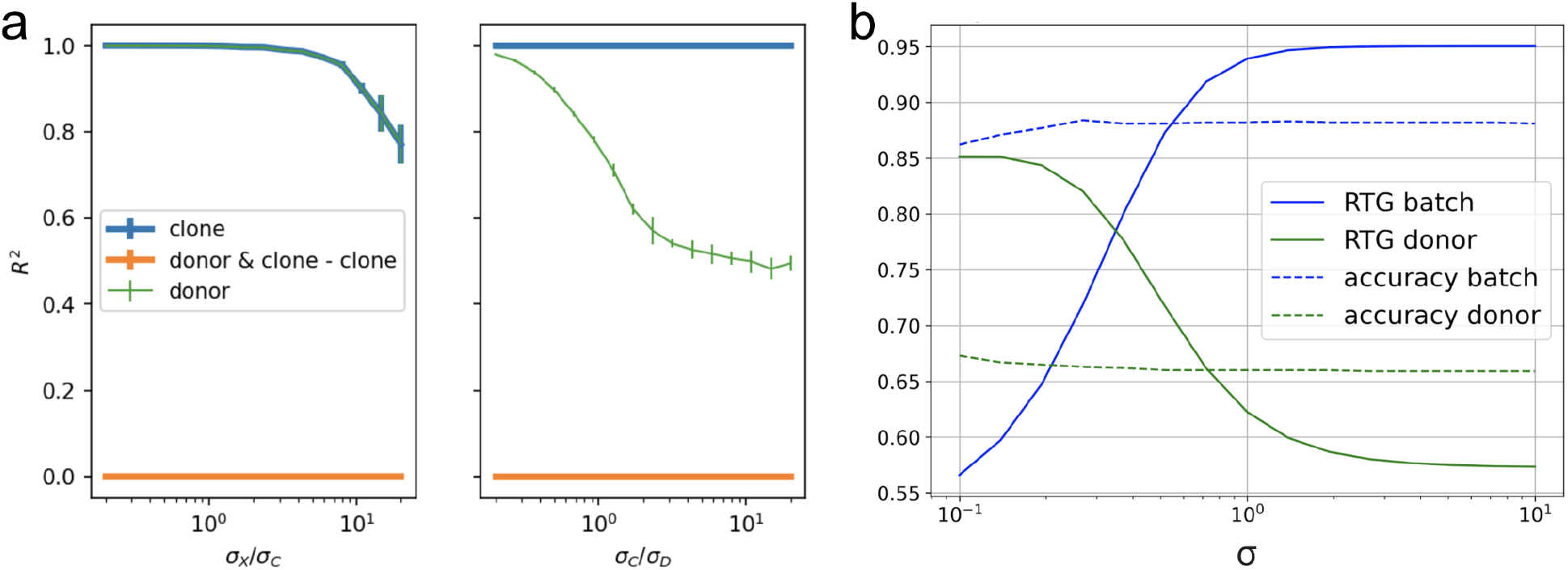
Weaknesses of encoder and decoder models for linear confounder discovery. **a**. Demonstration that linear isolation of donor confounding effects fails. As in Figure 2e, except for the orange line that attempts to isolate the confounding effects of donor by subtracting the variance explained by a linear model that regresses on clone alone from a model that regresses on both donor and clone. However, this quantity is alway zero because donor and clone are hierarchically nested confounders (i.e., all data samples that share the same clone label also share the same donor label), meaning that clone always provides perfect information about donor. Conventions as in Figure 2e except that σ_*D*_ = 40σ_*C*_ for the left panel. **b**. Linear decoders as confounder detectors are insensitive to effect size. A data generative process was used that could parametrically switch from having the variables donor or batch primarily structure the data. Decoding accuracy is insensitive to this switch while RTG scores report which variable is most important. Chance decoding accuracy is 0.1; differences between batch and donor decoding is due to finite size effects. 6-dimensional Gaussian mixture data was generated as follows. For dimensions *j* ∈ {1,2,3}, the donor and batch means were sampled as 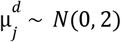 and 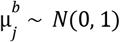. For dimensions *j* ∈ {4,5,6}, the means were sampled as 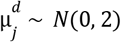 and 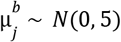. The number of donors and batches were each 100. Then, for the *i*^th^ of 1000 data points, a random donor *d_i_* and a random batch *b_i_* were chosen. For *j* ∈ {1,2,3}, the data point was sampled as 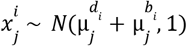, while for *j* ∈{4,5,6}, 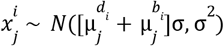.

**Supplemental Figure S2.**
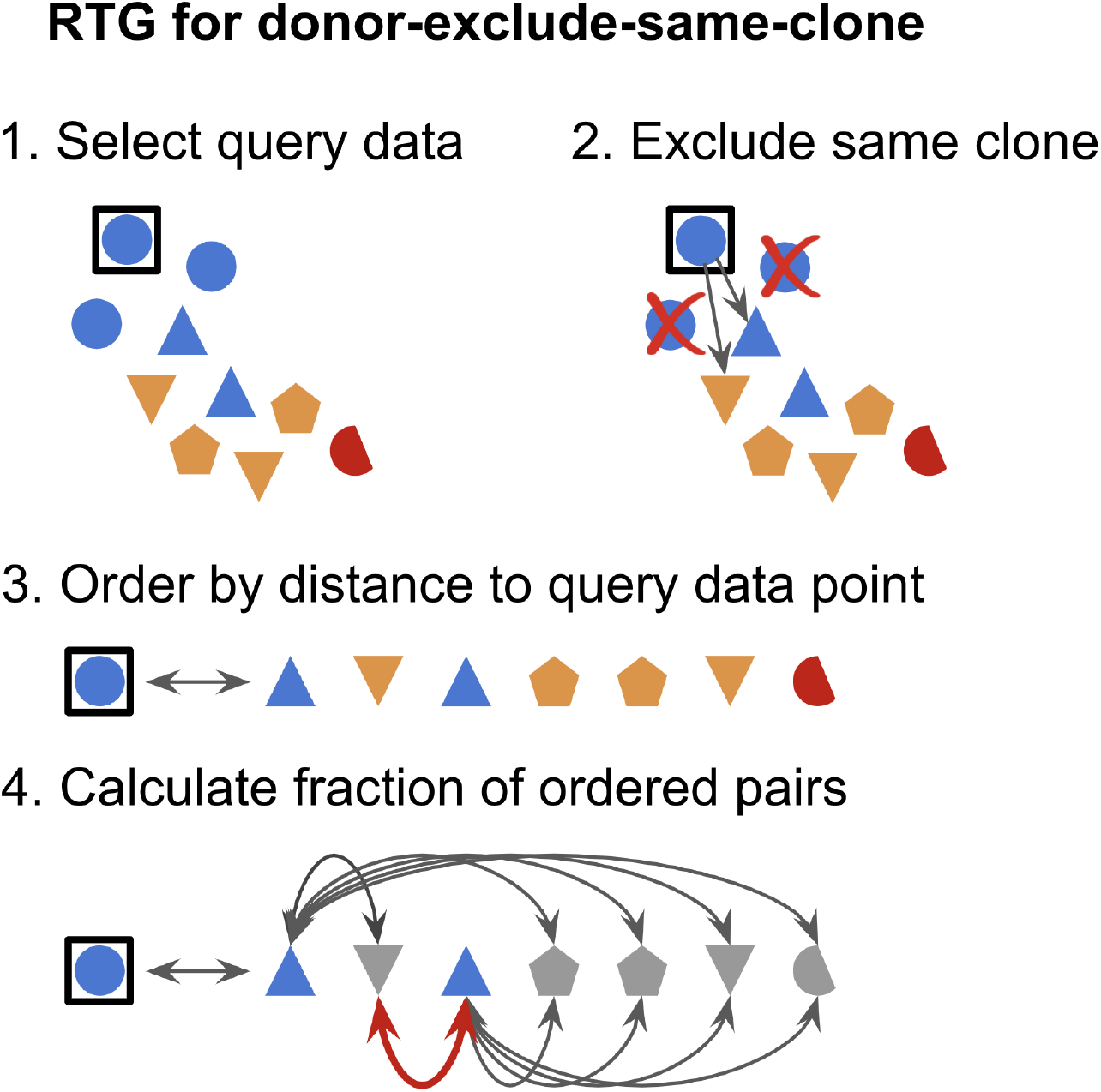
Diagrams illustrating the computation of the RTG query score for donor-exclude-same-clone (Algorithm 1 in Methods). First, a query data point is selected. Second, all data points that share the same clone as the query are excluded from the current iteration. Third, the data are ranked by their distances from the query. The arrows in diagram 2 show the distances from the query to the closest two non-excluded data points. Finally, when evaluating pairs of points – one that shares the same donor as the query and another that does not – the fraction of such pairs that are ordered by their distances to the query is the RTG query score for the current query. In diagram 4, one out of ten possible pairs is out of order (the pair marked with the red arrow), giving an RTG score for this query of 0.9.

**Supplemental Figure S3.**
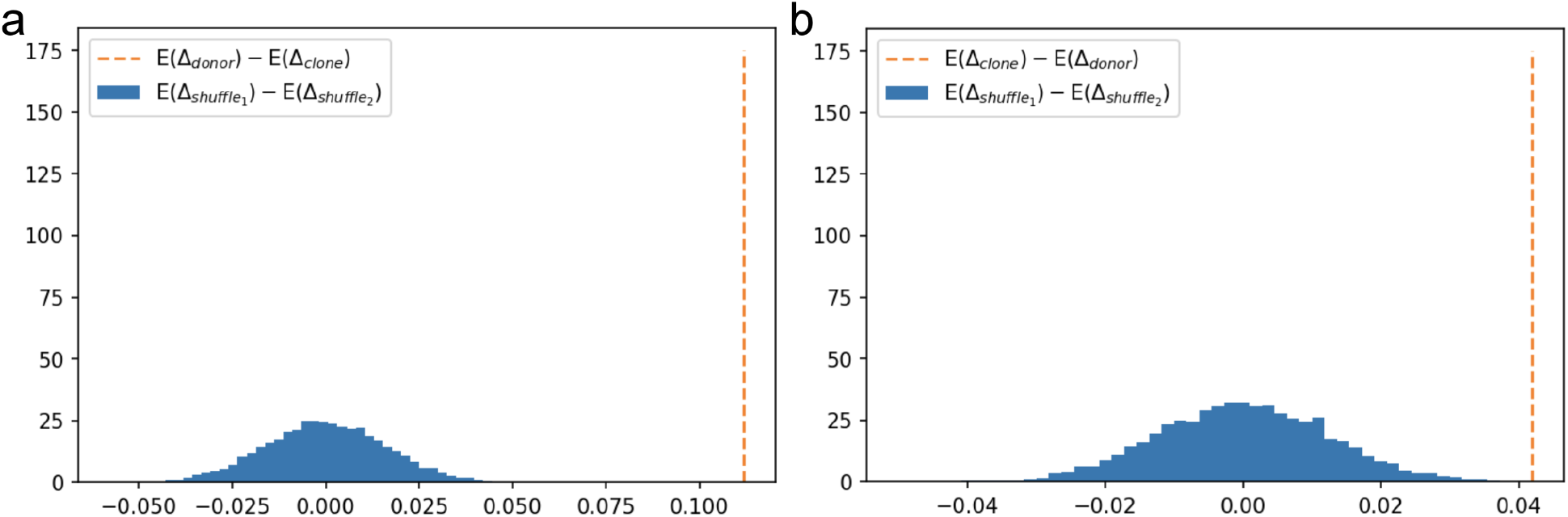
Shuffle distributions for changes from debiasing by donor versus clone. **a**. The mean difference in accuracies after and before debiasing by donor minus the mean difference in accuracies after and before debiasing by clone was calculated with the mean taken over 100 random samples of the data (dotted orange line). Then, the accuracy differences for donor and clone were shuffled 10000 times and for each shuffle the difference in mean differences was computed (blue histogram). No shuffles resulted in values greater than the orange line giving *p* < 10^−4^. **b**. Same as **a** with the roles of donor and clone reversed.

**Supplemental Figure S4:**
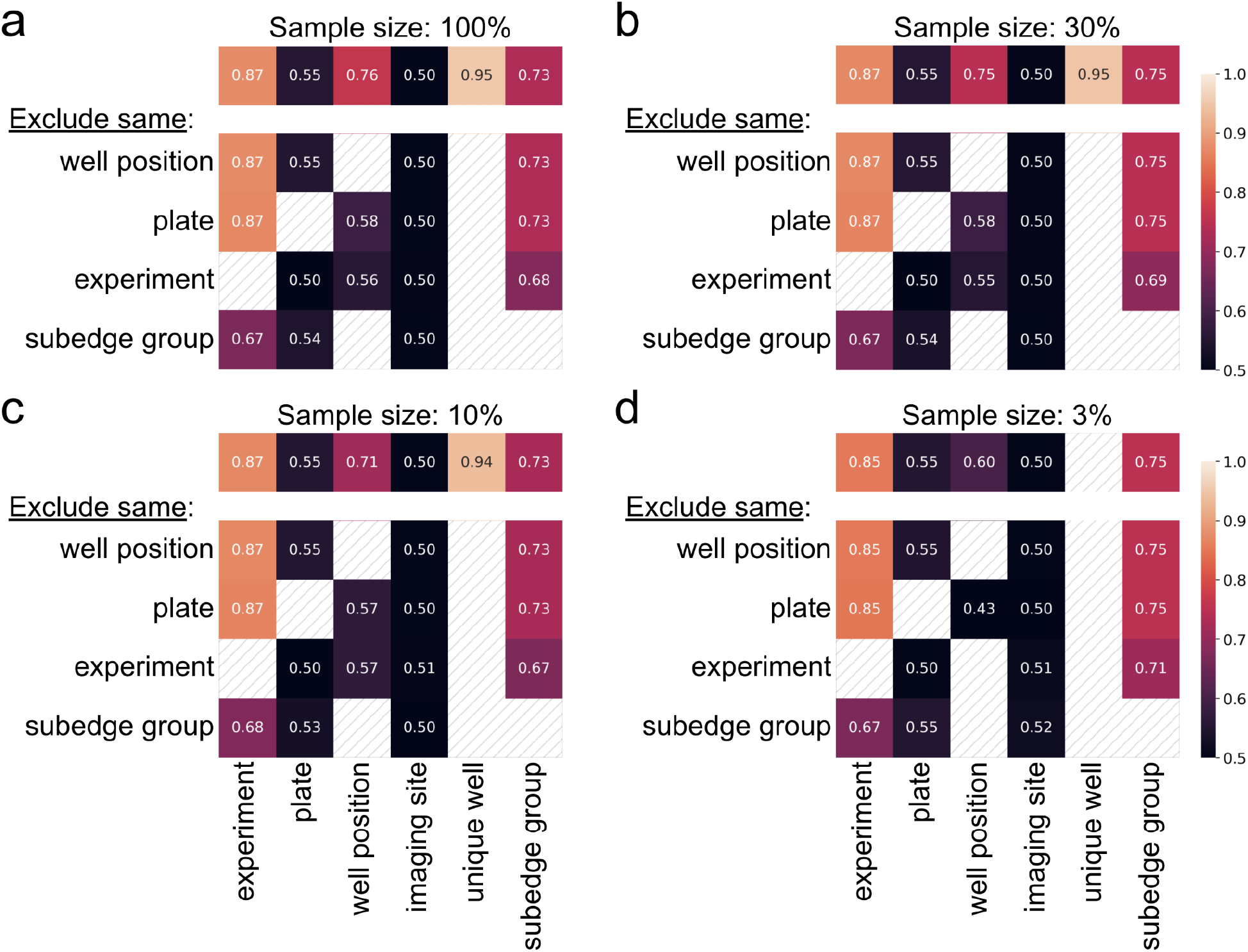
Comparing RTG scores between a full dataset from Recursion Pharmaceuticals (Cuccarese et al. 2020) and three subsampled versions of the same dataset. **a**. This panel is a repeat of Figure 5a. **b–d**. RTG scores after subsampling 30%, 10%, and 3% of the original data. Spearman rank correlations between the RTG scores before and after subsampling are 0.997, 0.985, and 0.879 for 30%, 10%, and 3% subsampling respectively.

